# Nuclear Progestin Receptor Mediated Linkage of Blood Coagulation and Ovulation

**DOI:** 10.1101/2021.11.08.467707

**Authors:** Jing Huang, Chao Sun, Dong Teng Liu, Nan Nan Zhao, Jordan A. Shavit, Yong Zhu, Shi Xi Chen

## Abstract

Ovulation is a dramatic remodeling process that includes rupture of blood capillaries and clotting, but coagulation is not thought to directly regulate this process. Herein, we report remarkable increases of coagulation factors V (*f5*, ∼3145-fold) and tissue factor (*f3a*, ∼120-fold) in zebrafish ovarian follicle cells during ovulation. This increase was mediated through the nuclear progestin receptor (Pgr), which is essential for ovulation in zebrafish, and was totally abolished in ovarian follicular cells from *pgr*^*-/-*^ mutants. In addition, promoter activities of *f5* and *f3a* were significantly enhanced by progestin (DHP) via Pgr. Similar regulation of human *F5* promoter activity was induced via human PGRB, suggesting a conserved mechanism. Site-directed mutagenesis of the zebrafish *f5* promoter further demonstrated a direct regulation of coagulation factors via progestin response elements. Moreover, a stark increase of erythrocytes occurred in capillaries meshed in wildtype preovulatory follicles but was absent in *pgr*^*-/-*^ mutants. Interestingly, anticoagulants significantly inhibited ovulation both *in vitro* and *in vivo*, respectively. Furthermore, reduced fecundity was observed in *f5*^*+/-*^ female zebrafish. Taken together, our study provides plausible evidence for steroid regulation of coagulation factors, and a new hypothesis for blood clotting triggered ovulation in vertebrates.

## 1. Introduction

Ovulation is a tissue remodeling process involving the rupture of follicular layers, basal membrane, and many blood capillaries intertwined in the follicular cells. This dramatic remodeling process shares several features with inflammatory responses (1). Blood coagulation was suggested to be part of the innate defense in response to inflammation during ovulation (2). The formation of blood clots at apical vessels was found shortly before follicle rupture in several animal species including sheep, rabbits, and rodents (1,3-5). Cessation of blood flow likely due to blood coagulation was also observed in apical vessels of ovarian follicles during the late ovulatory phase in women (6). In addition, an increase in activity and expression of thrombin (a protease essential for fibrin formation) and its receptors were also reported in luteinized granulosa cells and follicular fluid in human (7,8). Increased fibrinogen secretion in bovine granulosa cells was thought to be related to the proteolytic activity needed for follicle rupture. Circulating F3 (also known as tissue factor) was elevated in women with polycystic ovary syndrome that is characterized by chronic oligo-ovulation or anovulation (9). Although these reports imply a possible association between blood coagulation and the ovulation process, the mechanisms for this role in ovulation are still unclear.

Progestin and its nuclear progesterone receptor (PGR) are well-established initiators for vertebrate ovulation (10-15). Recently, we identified coagulation factors *f3a* and *f5* as potential downstream targets of Pgr in a transcriptomic analysis of preovulatory follicles from zebrafish (16). Interestingly, changes in these coagulation factors during ovulation appear to be conserved among human, mice, and zebrafish (16-18). In addition, the entire coagulation system and its factors are also highly conserved between zebrafish and mammals (19-21). Possible involvement of progestin or PGR in the regulation of coagulation have been hinted from several studies using progestin contraceptives. F3 was upregulated in endometriosis and at endometrial bleeding sites in women using long-term progestin-only contraception (22). Orally administered progesterone reduced spontaneous miscarriages and improved fertility among women who had a history of recurrent miscarriages due to abnormal bleeding in early pregnancy (23). Increased risk of venous thromboembolic events was associated with combined oral contraceptives, particularly those containing the progestin desogestrel (24,25). A recent genetic study indicated that present-day carriers of the Neandertal variant of PGR had less bleeding during early pregnancy because of high levels of PGR expression (26). However, evidence of progestin-induced or PGR-mediated regulation of coagulation factors, especially in ovarian follicles during ovulation, is lacking.

In this study, we examined the expression, hormonal regulation, and potential function of coagulation factors (*f5* & *f3*) in preovulatory follicles of zebrafish. The *f3* gene is duplicated in zebrafish (*f3a* and *f3b*) (27). We first demonstrated the expression and localization of *f5, f3a*, and *f3b* in preovulatory follicular cells. We found that progestin (DHP, 17α, 20β-dihydroxy-4-pregnen-3-one, a native ovulation-inducing ligand in zebrafish) via Pgr regulated *f5* and *f3a* expression in preovulatory follicles. We also found that follicular cells with strong Pgr expression were adjacent to blood capillaries, and inhibition of blood coagulation reduced ovulation. Finally, fertility and ovulation were reduced in *f5*^*+/-*^ zebrafish mutants *in vivo*. Our study provides credible evidence that coagulation factors (*f5* & *f3a*) in preovulatory follicular cells are regulated by progestin (DHP) via Pgr, which are well-known upstream initiators for ovulation. Our results also suggest a previously unknown role of these coagulation factors in ovulation and female fertility.

## 2. Materials and Methods

### 2.1 Animals & reagents

Zebrafish used are offspring of the Tübingen strain unless stated otherwise. Detailed information for knockouts, transgenic lines, key chemicals, and reagents are listed in Supplemental Table 1 & Table 2 (28). Fish were fed three times daily with a commercial food (Otohime B2; Marubeni Nisshin Feed, Tokyo, Japan), supplemented with newly hatched brine shrimp, and were maintained in recirculating system (ESEN, Beijing, China) at 28 ± 0.5 °C under a controlled photoperiod (lights on at 08:00, off at 22:00). The daily ovulation cycle in mature adult zebrafish is established according to an established protocol (29). Briefly, each mature adult female and male zebrafish were paired in a spawning tank. These fish typically start spawning soon after lights on in the morning. Around noon, fish water was replaced, and fish were fed at least three times daily with the commercial food and supplemented with newly hatched brine shrimp. Around one hour prior to lights off, fish water was replaced again, and a mesh inner tank was inserted into each spawning tank to prevent cannibalism typical for zebrafish adults. After one week, most fish form a daily ovulation cycle. When applicable, siblings from heterozygous in-cross were used for comparison. All experimental protocols were approved by the Institutional Animal Care and Use Committee (IACUC) at Xiamen University.

### 2.2 Collection of fully-grown follicles (stage IV) during a daily spawning cycle

Fully-grown immature follicles (stage IVa, >650 μm, with visible germinal vesicles, i.e., GV, appears opaque) or mature follicles (stage IVb, GV disappears, appears transparent) were collected at four representative timepoints during a daily spawning cycle (Fig. 1A): 12:00 (IVa follicles); 20:00 (IVa follicles); 05:00 (the onset of oocyte maturation, IVa follicles); and 06:40 (completion of oocyte maturation but prior to ovulation, IVb follicles). To be simple and accurate in our description, we will use “IVa” to indicate fully-grown immature stage IVa follicles; “IVb” to indicate mature but yet ovulated follicles (prior to ovulation) throughout this manuscript.

**Figure 1.**
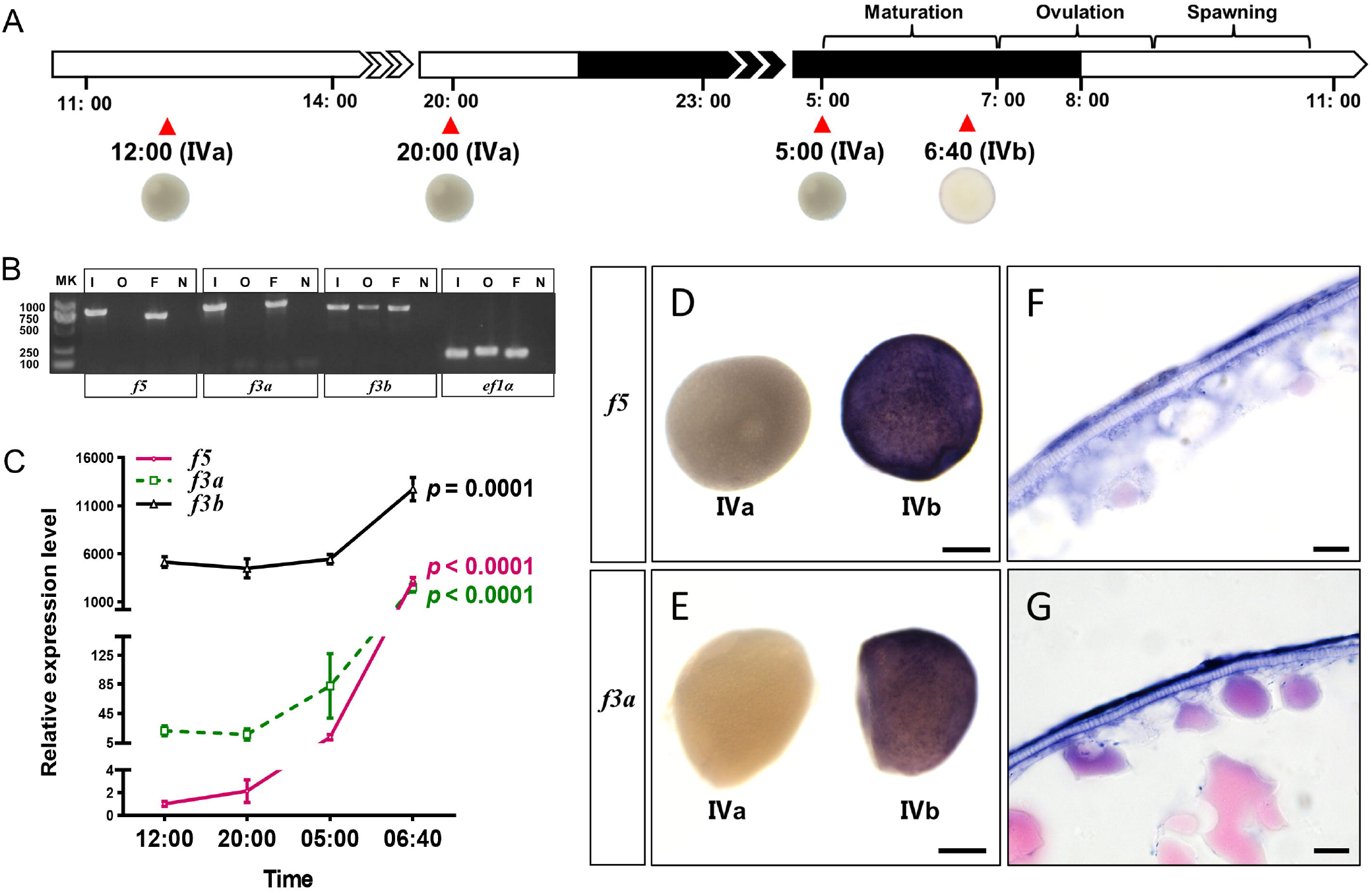
Dramatic increase of *f5* an*d f3a* transcripts in follicular cells of preovulatory follicles during a daily spawning cycle *in vivo*. (**A**) Schematic illustration of daily spawning cycle in zebrafish synchronized with lights on. Oocyte maturation in zebrafish typically occurs around 06:40, while ovulation occurs around 07:00. (**B**) Cell-specific expression of *f5, f3a, and f3b* in mature follicles examined by RT-PCR. Transcripts of *f5* and *f3a* were expressed in the follicular cells but not in the oocytes, while the transcript of *f3b* was expressed in both follicular cells and oocytes of mature preovulatory follicles. MK: molecular makers; I: intact follicles including both surrounding follicular cells and germ cell; O: denuded oocyte (without follicular cells); F: follicular cells (with no germ cell); N: no template control. (**C**) Dramatic increase of *f5, f3a*, and *f3b* transcripts in preovulatory follicles during a daily spawning cycle *in vivo*. Fully-grown ovarian follicles (stage IVa or IVb) were collected from wildtype mature female zebrafish at four representative timepoints during a daily spawning cycle (see Fig. 1A). Expression of *f5, f3a*, and *f3b* mRNA were determined by qPCR and normalized to the internal housekeeping gene (*ef1α*). Data are expressed as the mean ±SEM (N = 4) relative to the respective transcript levels of *f5* in IVa follicles collected at 12:00. P-values were calculated by one-tailed one-way ANOVA followed by Dunnett’s test against respective controls (samples collected at 12:00). (**D-G**) Dramatic increase of *f5* and *f3a* transcripts in the follicular cells of mature follicles. Fully-grown but immature (stage IVa) and mature follicles but prior to ovulation (stage IVb) were collected at 20:00 and 06:40, respectively. Strong signals of *f5* and *f3a* transcripts were only observed in mature follicles (IVb) by whole-mount *in situ* hybridization (WISH), while no signals were observed in the immature follicles (IVa, D, E). The strong ISH signals were localized in the follicular cells (F, G). Scale bars: 250 µm (D, E) and 10 μm (F, G).

To sacrifice fish humanely, the spinal cord and blood supply behind the gill cover were cut off swiftly using sharp scissors following a lethal dose of anesthetization (MS-222, 300 mg/L buffered solutions). Ovaries were immediately placed in 60 % L-15 media containing 15 mM HEPES at pH 7.2. Individual follicles were obtained according to an established procedure (29,30). At each timepoint, a total of 25 IVa or IVb follicles (containing both germ cell and surrounding follicular cells) from 4 females were pooled as one sample. In addition, the follicular cells (∼100 follicles) or their enclosed oocytes (∼20 follicles) were collected separately from IVb follicles according to an established method (31). These samples were immediately homogenized in RNAzol solution (Molecular Research Center, Cincinnati, USA) and stored in a -80 °C freezer until further processing.

### 2.3 Hormone treatments

According to the established follicle classification system (29), follicles at the different stages were collected at 14:00 from several females as described above. Intact follicles with no damage were transferred into a 24-well culture plate containing 60 % L-15 medium. These follicles were incubated at 28 °C with various doses of hormones or various exposure times. Several progestin concentrations (DHP, 1-1,000 nM) with a 2 hours (hrs) incubation or several exposure times (0.5-6 hrs) with 100 nM DHP were examined. Based on the result of dose-and time-dependent experiments, follicles at different stages were incubated with DHP (100 nM) for 2 hrs. Besides, stage IVa follicles were incubated with hCG (50 IU/mL) (30) or DHP (100 nM) alone or in combination for 2 hrs. Stage IVa follicles collected from *pgr*^*-/-*^ were incubated with DHP (100 nM) for 2 hrs. After incubation, minimal 25 follicles with no damage were collected and homogenized in RNAzol, stored in a -80 °C freezer until RNA purification. Minimal three replicates were collected at each timepoint.

### 2.4 Real-time quantitative PCR (qPCR)

RevertAid cDNA synthesis kit (Thermo Scientific, Waltham, USA) was used for reverse transcription (500 ng total RNA/sample). All qPCRs were carried out in a 20 μL reaction on the qTOWER 2.2 real-time PCR system (Analytik Jena AG, Jena, Germany) using the PowerUp SYBR green master mix (Applied Biosystems, USA). Each PCR primer of a pair was designed to target different exons of the gene (Supplemental Table 3) (28). Expression of the target genes was determined as relative values using the comparative C_t_ method (32) with the *ef1α* gene serving as an internal control.

### 2.5 Whole-mount *in situ* hybridization (WISH) and histological analysis

A DNA fragment corresponding to partial *f5* or *f3a* cDNAs was amplified from ovarian cDNAs by PCR using the gene-specific primers (see Supplemental Table 3 for detail) (28) and cloned into the pGEM-T Easy vector (Promega, USA). Correct sequence and orientations of inserts were validated by Sanger sequencing. Antisense or sense digoxigenin-labeled cRNA probes for *f3a* or *f5* were synthesized by PCR using the plasmid cDNA templates, SP6 or T7 RNA polymerase depending on the insert orientation. IVa or IVb follicles were collected at 20:00 and 06:40 (immediately prior to ovulation), respectively, and fixed in 4 % PFA overnight at 4 °C. Following washing with PBST, the follicles were treated with proteinase K (5 µg/mL) for 2 min at 37 °C, then washed and subsequently hybridized with digoxigenin-labeled *f5* or *f3a* sense or antisense probes at 65 °C overnight using a previously established protocol (33). Following WISH, whole follicles were photographed using an M165FC dissecting microscope and DFC 550 digital camera (Leica, Wetzlar, Germany). After imaging, follicles were dehydrated through increased ethanol concentration, embedded in paraffin, sectioned at 10 µm, mounted on a polylysine-treated glass slide, and stained lightly with eosin for better imaging. These sections of follicles were imaged using DM 2500 LED microscope and DFC 7000T digital camera (Leica).

### 2.6 Dual-luciferase reporter assay

The Pgr expression and reporter assay plasmid constructs were listed in Supplemental Table 4 (28). Site-directed mutagenesis of four potential Pgr response elements (PRE, i.e., Pgr binding sites) in zebrafish *f5* proximal promoter sequences (Fig. 4E, Supplemental Table 5) (28) was conducted using a Q5 site-directed mutagenesis kit (NEB, Ipswich, MA, USA) according to the manufacturer’s protocol. Human embryonic kidney cells (HEK 293T) were used for the transactivation assays (34,35). Briefly, cells were maintained at 37 °C with 5 % CO_2_ in phenol red-free Dulbecco’s modified eagle’s medium (Hyclone, USA) containing penicillin/streptomycin (Meilunbio, Dalian, China) and supplemented with 10 % fetal bovine serum (FBS, Gibco, Brazil). Lipofectamine 3000 (Life Technologies) was used for transient transfection according to manufactures’ protocols. Cells were transiently transfected with a firefly luciferase reporter plasmid (3 µg/vector/60mm petri dish), the pRL-TK vector (150 ng/60mm petri dish) containing the Renilla luciferase reporter gene (as an internal control for transfection efficiency), and a Pgr expression vector or a control vector with no insert (1.5 µg/vector/60mm petri dish). After an overnight incubation at 37 °C, the medium was replaced with luciferase assay medium (DMEM without phenol red, supplemented with 10 % charcoal-stripped FBS) containing various concentrations of progestins (100 pM-1 µM DHP for zebrafish Pgr, or P4 for human PGRB, respectively) or vehicle. Following additional 24 hrs of incubation at 37 °C, the cells were washed with PBS and harvested to determine firefly and Renilla luciferase activities according to the Dual-Luciferase reporter assay system (Promega, USA) on a luminometer (GloMax 20/20, Promega, USA). Firefly luciferase activities were normalized using Renilla luciferase data. After normalization for transfection efficiency, induction factors were calculated as the ratios of the average luciferase value of the steroid-treated samples versus vehicle-treated samples.

### 2.7 Blood capillaries in stage IV follicles

IVa immature follicles were obtained from paired female fish (*Tg(pgr:eGFP/fli1:DsRed)*). In this line, the blood capillaries are labeled with DsRed, while the follicular cells are labeled with eGFP. The nucleus of follicles was co-stained with Hoechst 33342 in a 29-mm glass-bottom dish (Cellvis, Hangzhou, China). Images were captured using an SP8 confocal laser scanning microscope (Leica, Germany). IVb mature follicles were obtained from ovaries undergoing ovulation from faired female fish (*Tg(fli1:eGFP)*) at 7:00. A series of images during ovulation was recorded using a Leica digital camera and fluorescent dissecting microscope described previously.

### 2.8 Erythrocyte numbers inside the capillaries on the surface of preovulatory follicles

Erythrocytes inside the capillaries on the surface of ovarian follicles were stained with *o*-dianisidine according to an established protocol (36). Ovaries with IVb follicles were collected at 6:40 or 7:00 from siblings of *pgr*^*+/-*^ in-cross, producing both wildtype (*pgr*^*+/+*^) and *pgr*^*-/-*^. As sampling moves closely toward 7:00, IVb follicles will be further in the ovulation process in *pgr*^*+/+*^ females. In contrast, IVb follicles in *pgr*^*-/-*^ females will never initiate ovulation (13,15), which serves as a control. After collection, the whole ovaries were immediately stained for 15 min at room temperature in the dark in 0.6 mg/ml *o*-dianisidine, 0.01 M sodium acetate (pH 4.5), 0.65 % H_2_0_2_, and 40 % (vol/vol) ethanol. Stained ovaries were washed 3 times for 2 min each with PBST. At least 3 ovary samples were collected at each timepoint for each genotype. Minimal 10 IVb follicles were randomly chosen from each sample for image capture using DM 2500 LED microscope and DFC 7000T digital camera (Leica), then the number of erythrocytes under the field of view was counted by a reviewer who was blinded from sampling conditions and genotypes.

### 2.9 *In vitro* ovulation assay

Ovaries with IVb mature follicles from *Tg(fli1:eGFP)* zebrafish were collected at 06:40, 20 min before ovulation, and were placed in 60 % L-15 medium. Each ovary was immediately cut into three equal pieces and then transferred into medium containing vehicle, 2 mM EDTA, or an anticoagulant (100 µg/mL heparin, 50 µg/mL warfarin, 50 µM dabigatran etexilate, or 250 µM rivaroxaban), to examine the inhibitory effect of anticoagulants on ovulation. The dose of each drug was chosen based on previous reports (37-39). Follicles were immediately dispersed by pipetting using a glass pipette (∼ 1 mm in diameter). At least 50 IVb preovulatory follicles per well were used in each experiment. The follicles were cultured at 28 °C in a 2 mL culture medium for 2 hrs. Ovulation was continuously monitored until 09:00 under a fluorescent dissecting microscope (Leica M165FC), and the numbers of oocytes that had successfully ovulated were recorded. Minimal replicates for each treatment were seven times.

### 2.10 *In vivo* ovulation assay

We chose warfarin over three other available anticoagulants, mainly because heparin has pleiotropic effects, and there are well-established and specific anticoagulant effects of warfarin on hepatocytes, major cells for producing coagulation factors for the systemic circulation. We added warfarin directly to fish tank water at a final concentration of 5 µg/ml (40) for examination of whether inhibition of coagulation would prevent ovulation. Fertile females were treated continuously with warfarin for 4 days with replacements of fresh water and warfarin every 12 hrs (40). For the ovulation/spawning test, each warfarin-treated female was paired with a fertile male in a spawning tank containing warfarin (final concentration, 5 µg/ml) in the evening (∼21:00) before the spawning test next day. Simultaneously, each control female (non-exposure) was paired with a fertile male in a spawning tank without warfarin. A blinded observer recorded the occurrence of ovulation/spawning by counting total and fertilized embryos at noon the following day. In the first trial, 34 females were randomly divided into either warfarin-treated or control (non-exposure) group. In the second trial, females from the control group of the first trial were randomly divided into either as control or warfarin treated. In the third trial, 42 females from a new batch were randomly assigned as warfarin treated or control group. 12 females from each group were randomly selected to examine the occurrence of ovulation/spawning. The remaining females from each group were randomly selected for collection of mature ovarian follicles (stage IVb) at 07:00 and erythrocytes were visualized using o-dianisidine staining (see section 2.8). Minimal six mature follicles were randomly chosen from at least three warfarin-treated or three non-treated female zebrafish. The number of erythrocytes in capillaries were counted by a blinded observer.

### 2.11 Consecutive spawning test

The spawning of zebrafish was optimized under enhanced feeding conditions as described previously (29). Multiple pairs and spawning tanks were set up. Ten to twenty mature females (three months old) of *f5*^*+/-*^ or wildtype (*f5*^*+/+*^) female siblings were used. Each female was individually paired and housed with a fertile wildtype male. Around 21:00 (1 hrs prior to lights off), water in spawning tanks was replaced, and a mesh inner tank was inserted into each spawning tank to prevent the eating of eggs by the adults. The number of fertilized eggs was recorded for each pair of fish around noon of the next day. Then, water would be replaced, and fish would be fed at least three times a day. The *f5*^*+/-*^ and wildtype (*f5*^*+/+*^) females were continuously monitored for at least 3 weeks. Data from the last two weeks were used for analyses.

### 2.12 Statistical analyses

Data analyses were performed using GraphPad Prism version 8. Depending on the experimental setup, Student’s t-test (unpaired or paired), or one-way ANOVA (one-tailed) followed by the Dunnett’s post hoc test was used to assess statistical differences in comparison to the control group. Each experiment (e.g. qPCR, reporter assay, WISH, drug exposure, reporter assay, erythrocytes count, *in vitro* ovulation, fertility test) was repeated at least once.

## 3. Results

### 3.1 Dramatic increase of *f5* an*d f3a* transcripts in follicular cells of preovulatory follicles during a natural spawning cycle *in vivo*

Expression of *f5* and *f3a* mRNA was exclusively restricted to the follicular cells of mature follicles (IVb) and was not found in the germ cells (oocytes) (Fig. 1B). In contrast, *f3b* mRNA was expressed in both follicular cells and oocytes (Fig. 1B). The expression of *f5, f3a*, and *f3b* in mature follicles (IVb), just prior to ovulation, collected at 06:40 were ∼3,145-fold, ∼120-fold, and ∼2-fold higher than those in immature but fully-grown follicles (IVa, collected at 12:00, Fig. 1C). Since the increase in *f3b* expression was modest, we focused the rest of our efforts on *f5* and *f3a*. Results from the whole-mount *in situ* hybridization (WISH) showed that both *f5* and *f3a* were nearly undetectable in immature follicles (IVa, Fig. 1D, E). In contrast, strong signals of both *f5* and *f3a* transcripts were detected in the mature follicles (IVb, Fig. 1D, E), while no signal was detectable using sense probes (data not shown). Sections of the mature follicle (IVb) from WISH showed strong signals of both *f5* and *f3a* transcripts in the follicular cells, while weak signals were observed in the chorion, and no signal was observed inside the oocytes (Fig. 1F, G).

### 3.2 Progestin upregulates *f5* and *f3a* transcripts through Pgr

#### 3.2a Transcripts of *f5* and *f3a* were primarily upregulated by progestin in preovulatory follicles

The progestin (DHP) is a well-established native ligand for Pgr and an upstream hormonal inducer for ovulation in zebrafish (13,34,41). As predicted, the increase of *f5* and *f3a* transcripts were dose- and time-dependent in fully grown immature follicles (IVa) exposed to external progestin (Fig. 2A-D).

**Figure 2.**
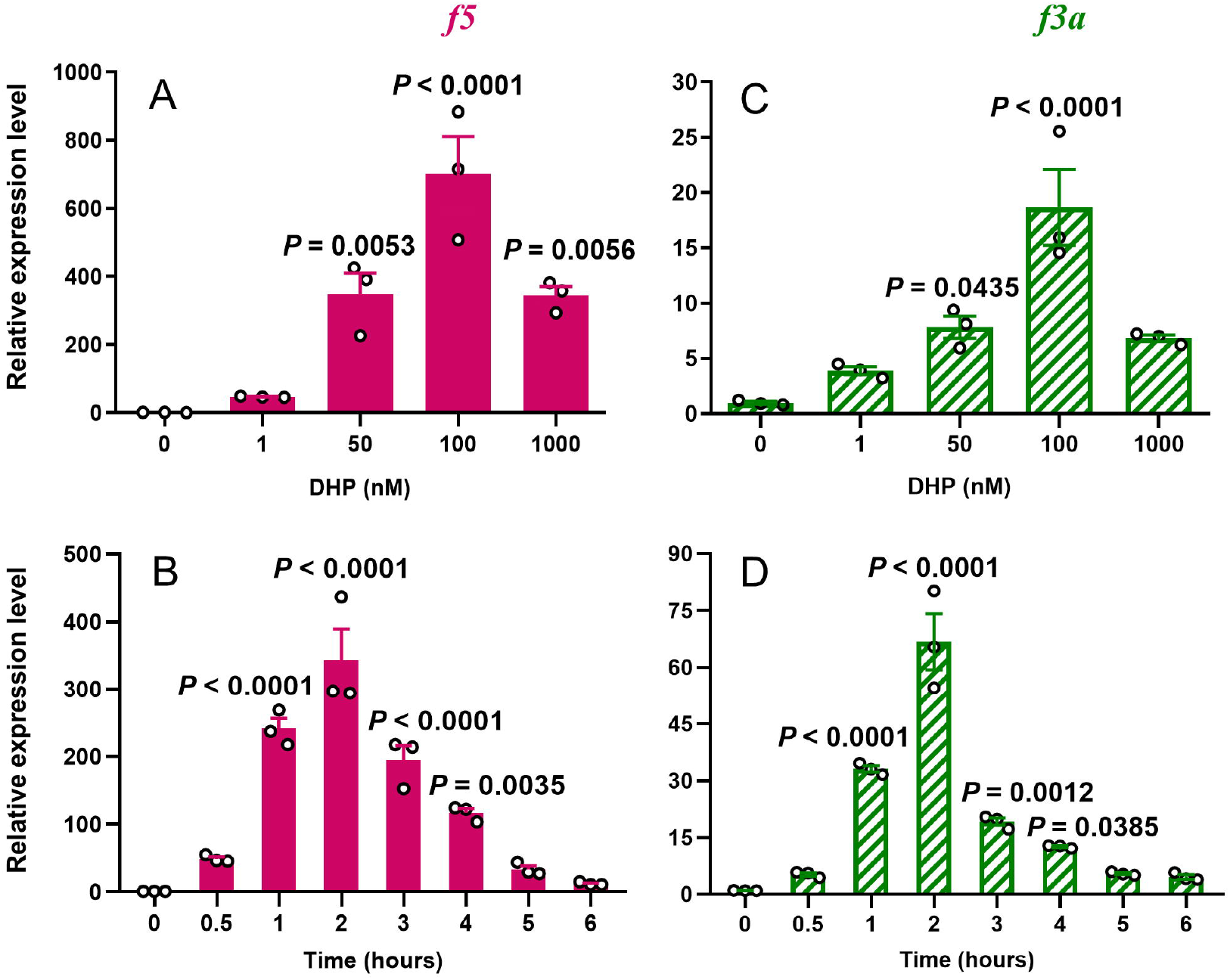
Transient increase of *f5* and *f3a* expression in preovulatory follicles in response to stimulation of progestin *in vitro*. (**A, C**) Effects of various doses of progestin (DHP) (2 hours exposure) on *f5* and *f3a* expression in fully grown immature preovulatory follicles (stage IVa) *in vitro*. (**B, D**) Effects of DHP (100 nM) on *f5* and *f3a* expression in preovulatory follicles (IVa) at various timepoints *in vitro*. P-values were calculated by one-tailed one-way ANOVA followed by Dunnett’s test against respective controls (0 timepoint or vehicle control).

Progestin increased *f5* expression in vitellogenesis follicles (III, ∼7 fold) and fully-grown immature follicles (IVa, ∼337 fold) (Fig. 3A). In comparison, only fully-grown immature follicles (IVa) responded to progestin with an increase in *f3a* transcripts, but not the early developing follicles (stage I-III, Fig. 3B). LH is the primary endocrine factor for inducing ovulation in all animals including zebrafish (42). As expected, increased *f5* expression (∼4 fold) was observed in IVa immature follicles exposed to hCG (50 IU/mL, Fig. 3C), but not for *f3a* (Fig. 3D). In comparison, progestin (DHP) had much stronger stimulatory effects on the expression of *f5* (∼318 fold) and *f3a* (∼10 fold) than hCG (Fig. 3C, D).

**Figure 3.**
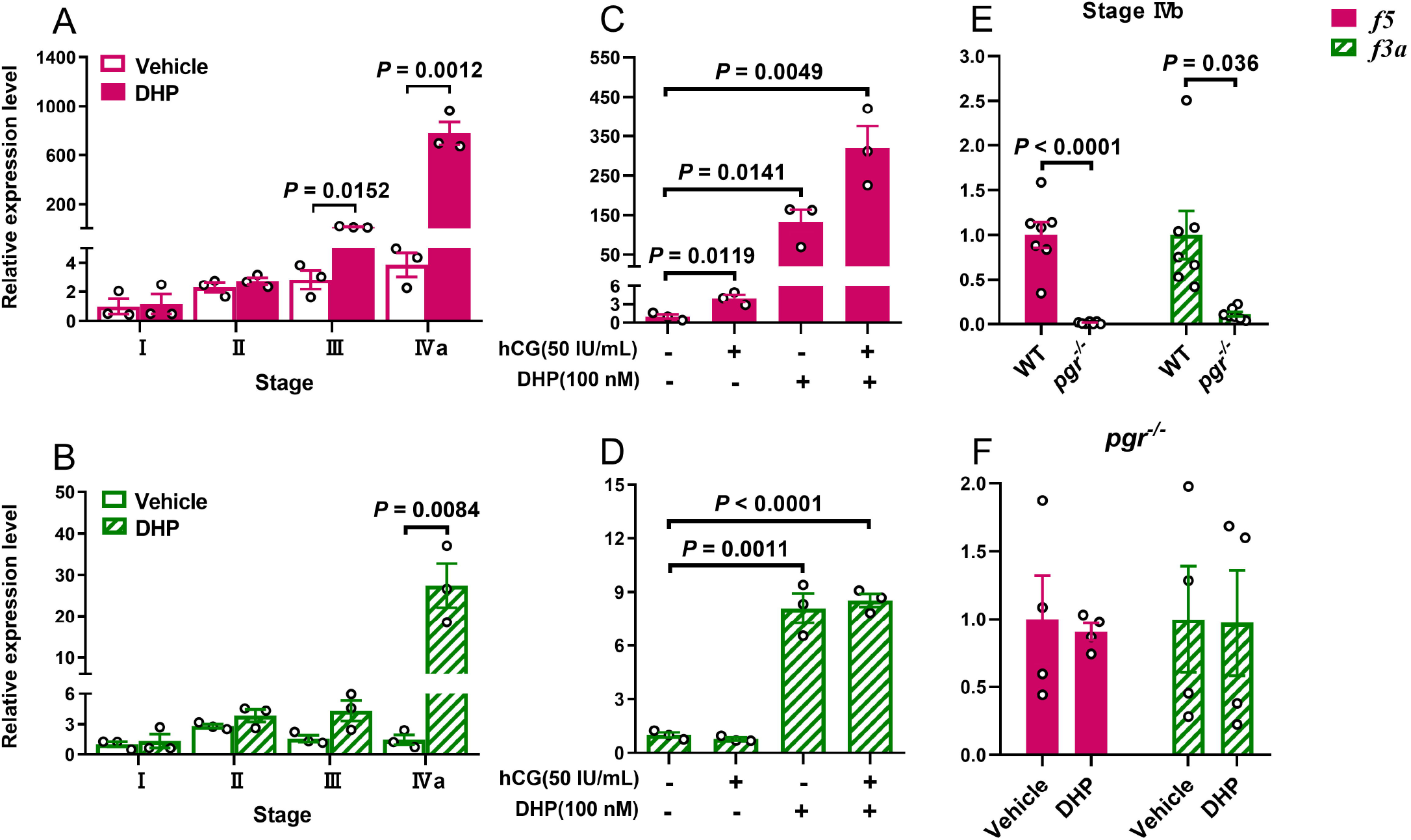
Increase of *f5* and *f3a* transcripts in late-stage follicles is mediated by Pgr. Follicles were incubated with progestin (DHP, 100nM), or LH (hCG, 50 IU/ml) alone, or in combination for 2 hrs at 28 °C *in vitro* before qPCR analysis. (**A**) An increase of *f5* transcript was observed in late**-**stage (III and IVa) follicles from wildtype (*pgr*^*+/+*^) zebrafish, but not in early-stage follicles (I or II) from wildtype zebrafish, in response to progestin stimulation. (**B**) An increase of *f*3a expression was observed only in fully-grown immature follicles (IVa) from wildtype (*pgr*^*+/+*^) zebrafish, but not in developing follicles (I-III) from wildtype fish, in response to progestin stimulation. (**C**) Synergistic increase of *f5* mRNA by hCG and DHP in fully grown immature follicles (IVa) from wildtype zebrafish. (**D**) Progestin but not hCG stimulated the expression of *f3a* mRNA in full-grown immature follicles (IVa) from wildtype zebrafish. (**E**) Both *f3a* and *f5* transcripts in preovulatory follicles (stage IVb) of *pgr*^*-/-*^ were significantly lower than that of *pgr*^*+/+*^ (WT). (**F**) Stimulatory effect of progestin (DHP) for *f3a* and *f5* expression was abolished in the stage IVa follicles of *pgr*^*-/-*^ mutants. The expression of *f5* and *f3a* transcripts were determined by qPCR and normalized to an internal control (*ef1α*). Data are expressed as the mean ± SEM (N = 3) relative to the respective transcript levels of target gene measured in wildtype, or vehicle treatment group. P-values were calculated by two-tailed Students’ t-test.

Significantly reduced *f5 and f3a* expression was found in mature preovulatory follicles (IVb) collected from *pgr*^*-/-*^ fish compared to those in wildtype siblings (Fig. 3E). Importantly, DHP-induced *f5* and *f3a* expression were blocked in fully-grown immature follicles (IVa) from *pgr*^*-/-*^ fish (Fig. 3F).

#### 3.2b Progestin modulates promoter activities of *f5* and *f3a* via Pgr

To further examine whether *f5* or *f3a* has PRE and whether progestin can directly activate these PREs via Pgr, we downloaded upstream sequences of *f5* and *f3a* of zebrafish, mouse, and human from the Ensembl genome database (https://asia.ensembl.org/index.html). Potential PREs were cross referenced and identified using several search engines and transcription factor databases (http://bioinfo.life.hust.edu.cn/hTFtarget#!/, TRANSFAC, JASPAR, CIS-BP, and HOCOMOCO, http://bioinfo.life.hust.edu.cn/HumanTFDB/#!/tfbs_predict; Supplemental Tables 5 & 6). We identified four potential PREs upstream (−2063/+32, ENSDARG00000055705) of the zebrafish *f5* gene, 12 putative PREs upstream (−2601/+202, ENSG00000198734) of the human *F5* gene, and 22 presumed PREs upstream (−2712/+125, ENSMUSG00000026579) of the mouse *F5* gene (Supplemental Table 5). We also detected one potential PRE upstream (−1892/+133, ENSDARG00000099124) of the zebrafish *f3a* gene, five putative PREs upstream (−2712/+123, ENSG00000117525) of the human *F3* gene, and 21 PREs upstream (−2640/+178, ENSMUSG00000028128) of the mouse *F3* gene (Supplemental Table 6).

Thereafter, we employed a dual-luciferase reporter assay to further verify our predictions with a focus on zebrafish f5 (*zf5*), zebrafish f3a (*zf3a*), and human F5 (*hF5*, Fig.4). A reporter vector containing MMTV (mouse mammary tumor virus) promoter with known PREs was used as control (Fig. 4A) (43). In the presence of Pgr, DHP, a native ligand for zebrafish Pgr, significantly increased the *f5 and f3a* promoter activities, and this was dose-dependent (Fig. 4B, C). No change in promoter activity was observed when pcDNA3.1(+)-Pgr was replaced by a control vector (pcDNA3.1(+), Fig. 4B, C). The increase for *f5* was much larger than *f3a*. In the presence of human PGRB, progesterone (P4), the native ligand of human PGRB, also significantly increased human *F5* promoter activity in a dose-dependent manner (Fig. 4D). No change in promoter activity was observed when pCMV-HA-hPGRB was replaced by a control vector (pCMV-HA, Fig. 4D). Because PGR can mediate transcriptional activation of genes that lack a canonical PRE or acts indirectly by binding to other co-factors (44), we further examined these four putative PREs in the *f5* promoter region using site-directed mutagenesis (Supplemental Table 5) that significantly decreased zebrafish *f5* promoter activity (Fig. 4E) (28). A PRE (5’-TAAAAAATGTCCT-3’) localizing to -1438 to -1425 of zebrafish *f5* appears to be the primary mediator of Pgr regulation (Fig. 4E).

**Figure 4.**
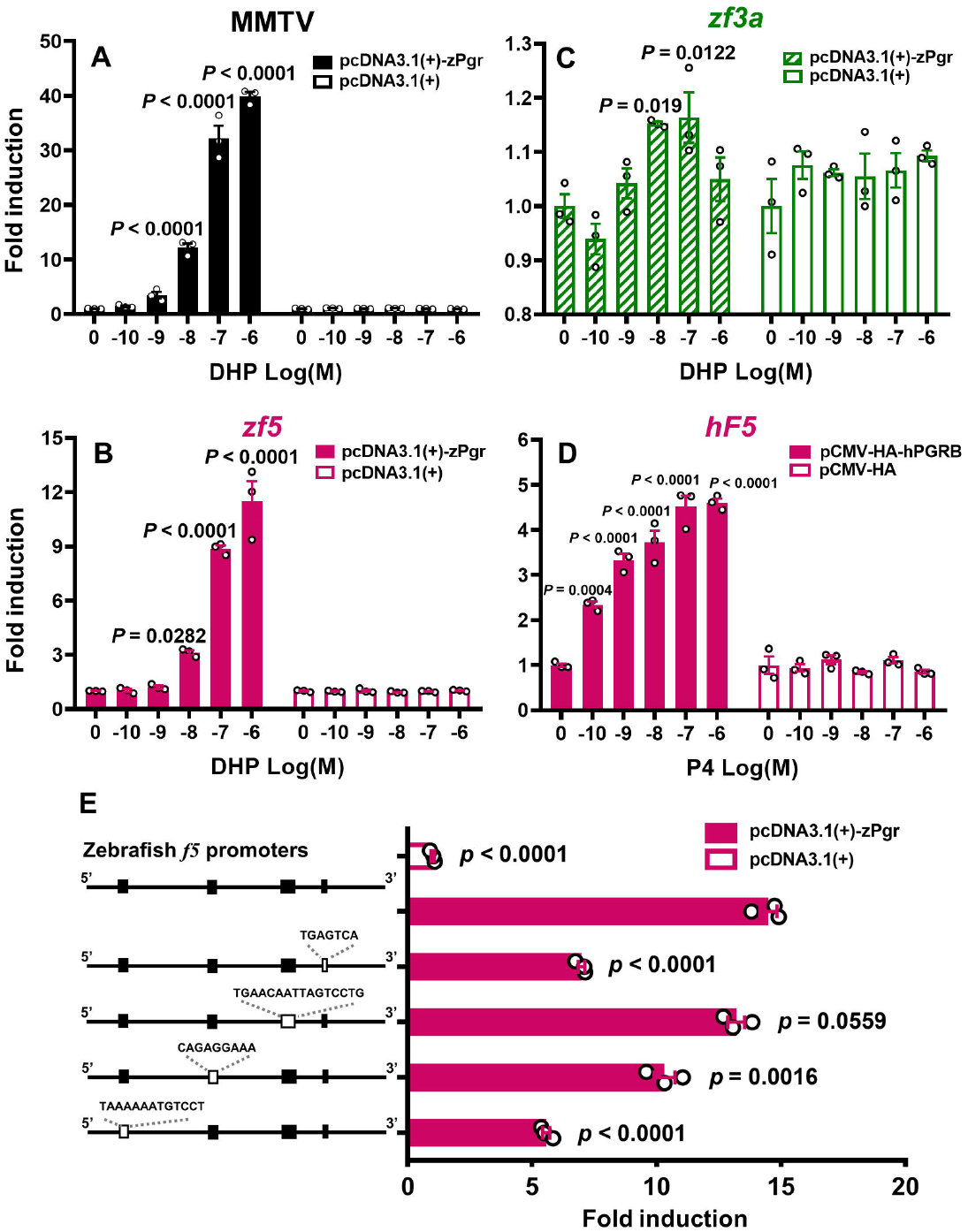
Progestin enhances promoter activity of *f5* and *f3a* via Pgr. HEK293T cells were transiently co-transfected with a firefly luciferase reporter vector containing putative Pgr binding elements (3 μg/vector/60mm petri dish), a pRL-TK vector (150 ng/60mm petri dish) containing the Renilla luciferase reporter gene (as a control for transfection efficiency), and a Pgr expression vector (1.5 μg/vector/60mm petri dish) or an empty control expression vector (1.5 μg/vector/60mm petri dish). In the presence of zebrafish Pgr (zPgr), progestin significantly increased MMTV (**A**), *zf5* (**B**), and *zf3a* (**C**) promoter activities. This increase was dose-dependent. In the presence of human PGRB (hPGRB), P4 also significantly increased human *hF5* promoter activity, and the increase was dose-dependent (**D**). Site-directed mutagenesis of four likely progestin receptor response elements (PRE) decreased the *zf5* promoter activity (**E**). HEK293T cells were incubated for 24 hrs with increasing concentrations of progestins (DHP or P4: 100 pM to 1 µM). Extracts of the treated cells were assayed for luciferase activity. Values are shown relative to the luciferase activity of the vehicle treatment group. Data are expressed as mean ± SEM (N = 3). P -values were calculated by one-tailed one-way ANOVA followed by Dunnett’s test against respective controls (vehicle controls, zero dose).

### 3.3 Coagulation factors contribution to ovulation

#### 3.3a Follicular cells with strong Pgr expression localize adjacent to blood vessels

We generated a double transgenic line *Tg(pgr:eGFP/fli1:DsRed)*, in which vascular endothelial cells are labeled by red fluorescent proteins, while *pgr* promoters drive strong GFP signals in Pgr expressing cells. Intriguingly, we found almost all the follicular cells with strong Pgr expression were located adjacent to capillary vessel networks on the surface of the IVa follicles in this dual transgenic zebrafish line *Tg(pgr:eGFP/fli1:DsRed)* (Fig. 5).

**Figure 5.**
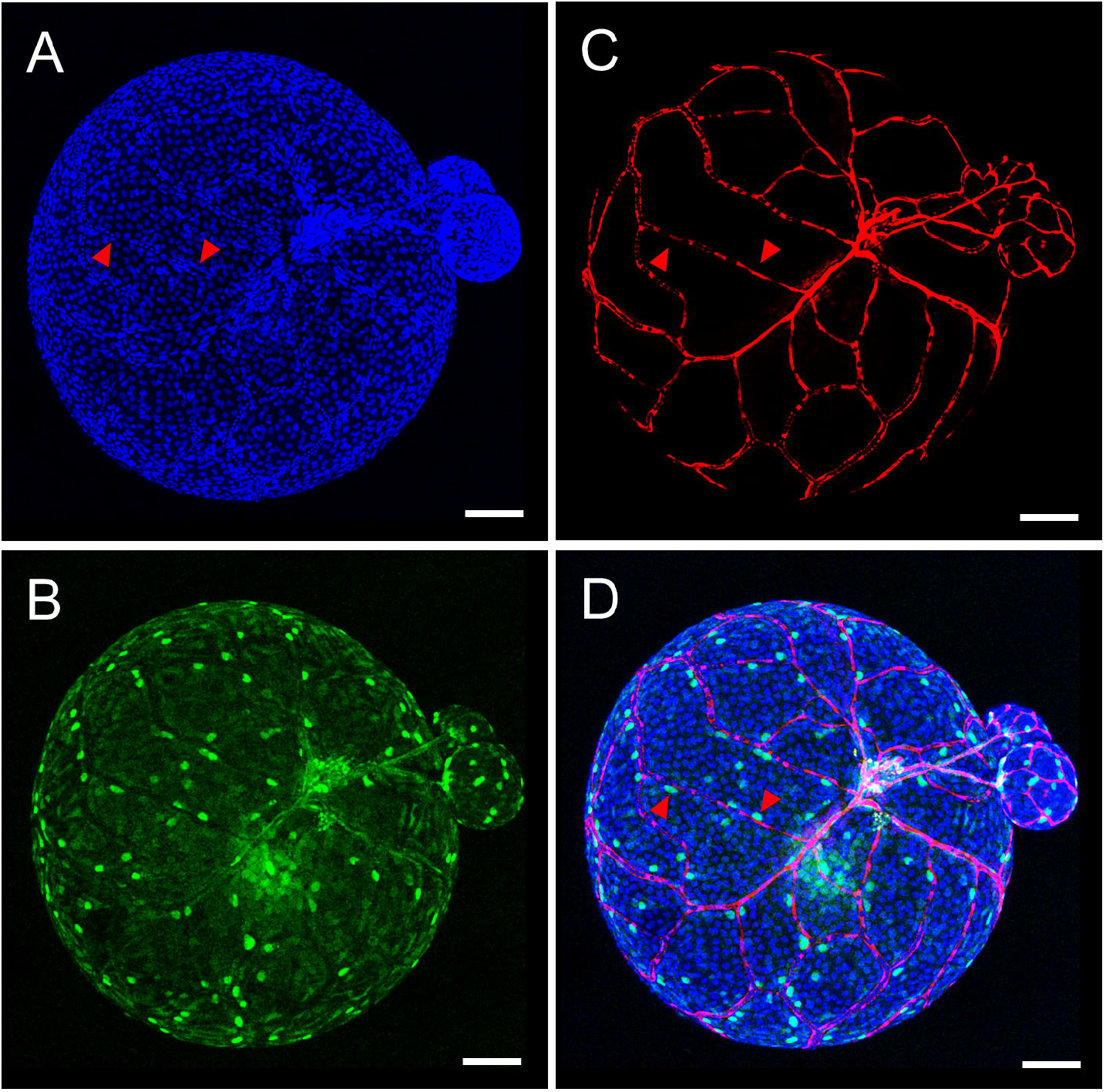
Follicular cells with strong Pgr expression are adjacent to blood capillaries. Images of a representative stage IVa follicle from a double transgenic zebrafish line *Tg(pgr:eGFP/fli1:DsRed)* are shown. (**A**) Nucleus staining of follicular cells by Hoechst 33342. (**B**) Pgr promoter drives GFP expression in follicular cells. (**C**) Red fluorescence-labeled vascular endothelial cells on the follicles. (**D**) Merged images to show that follicular cells with strong GFP expression driven by the Pgr promoters are adjacent to blood capillaries. Red arrowheads indicate follicular cells with strong GFP expression driven by the Pgr promoters. Scale bars: 100 μm.

#### 3.3b Rupture of blood vessels during ovulation

The ovulation process started with the appearance of a small hole in follicular cells and a ruptured blood vessel (indicated by a red arrow in Fig. 6A, supplemental video clip A) (28) with a concomitant appearance of the ovulating oocyte on the surface of the ovaries. When the hole of the broken follicular cell layer reached a certain size, the oocyte was quickly released from the follicular layer (Fig. 6B-E, supplemental video clip A) (28).

**Figure 6.**
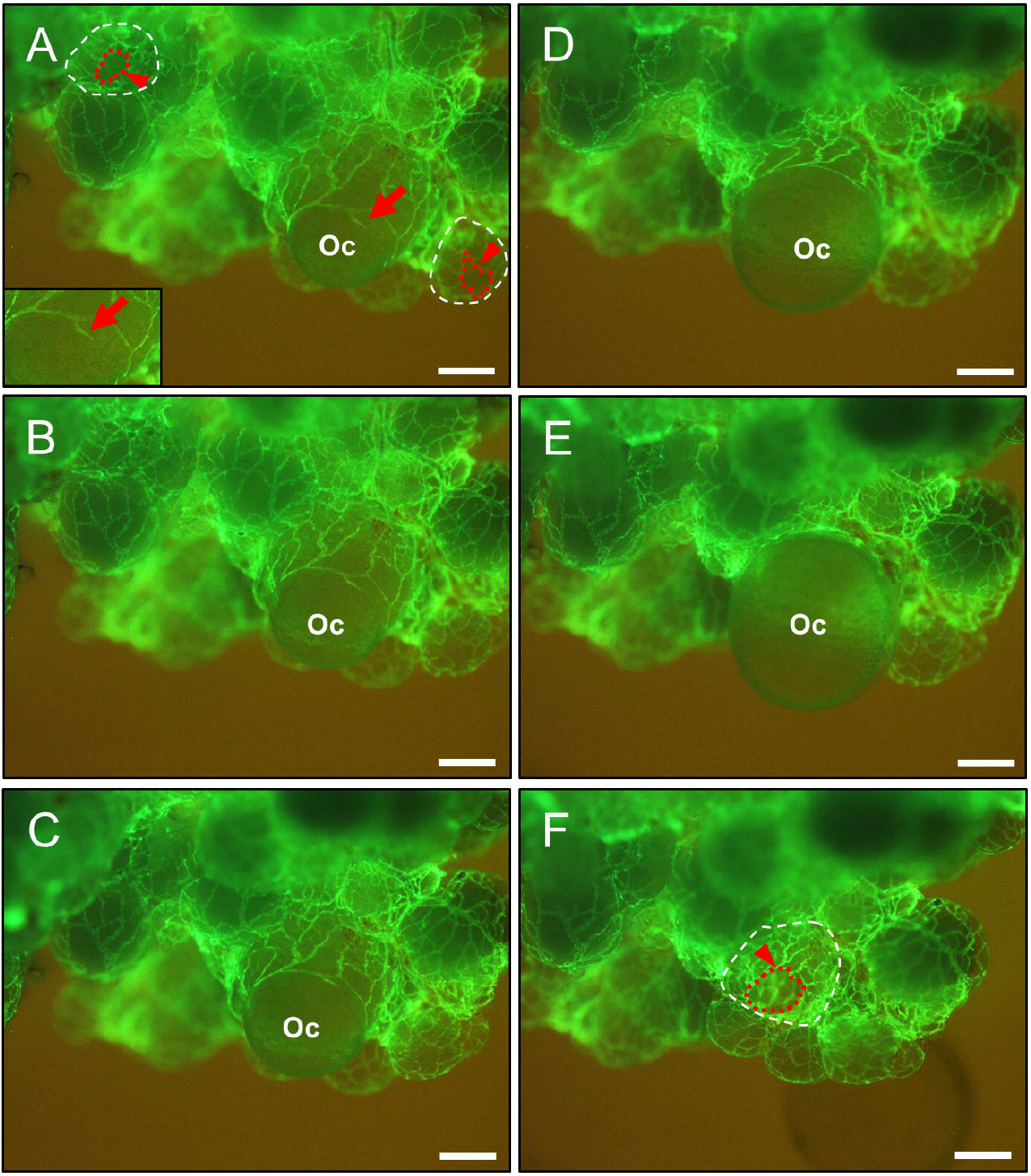
Rupture of blood capillaries and follicular cells during ovulation. Ovaries undergoing ovulation were collected from *Tg(fli1:eGFP)* females at 07:00. (**A-F**), a series of images were taken when a mature (stage IVb) ovarian follicle started ovulation. Interconnected blood vessels labeled with GFP were observed easily on the surface of ovarian follicles. A red arrow indicates a ruptured blood vessel on the surface of an ovulating follicle. Red arrowheads and red dotted lines indicate a hole left in the remaining follicular cells after ovulation. The white dotted line represents the remaining follicular cells. Scale bars: 250 μm. Oc, oocyte. (See supplemental video clip A for additional details) (28).

#### 3.3c Increased numbers of erythrocytes appear in the lumen of blood vessels of mature preovulatory follicles as ovulation proceeds

The numbers of erythrocytes in capillaries of mature preovulatory follicles (IVb) increased as they proceeded to follicular rupture, from ∼06:40, (Fig. 7A, D, G) to ∼07:00 (Fig. 7B, E, G). This effect was absent in the same mature follicles (IVb) collected from anovulatory *pgr*^*-/-*^ female siblings around 07:00 (Fig. 7C, F, G).

**Figure 7.**
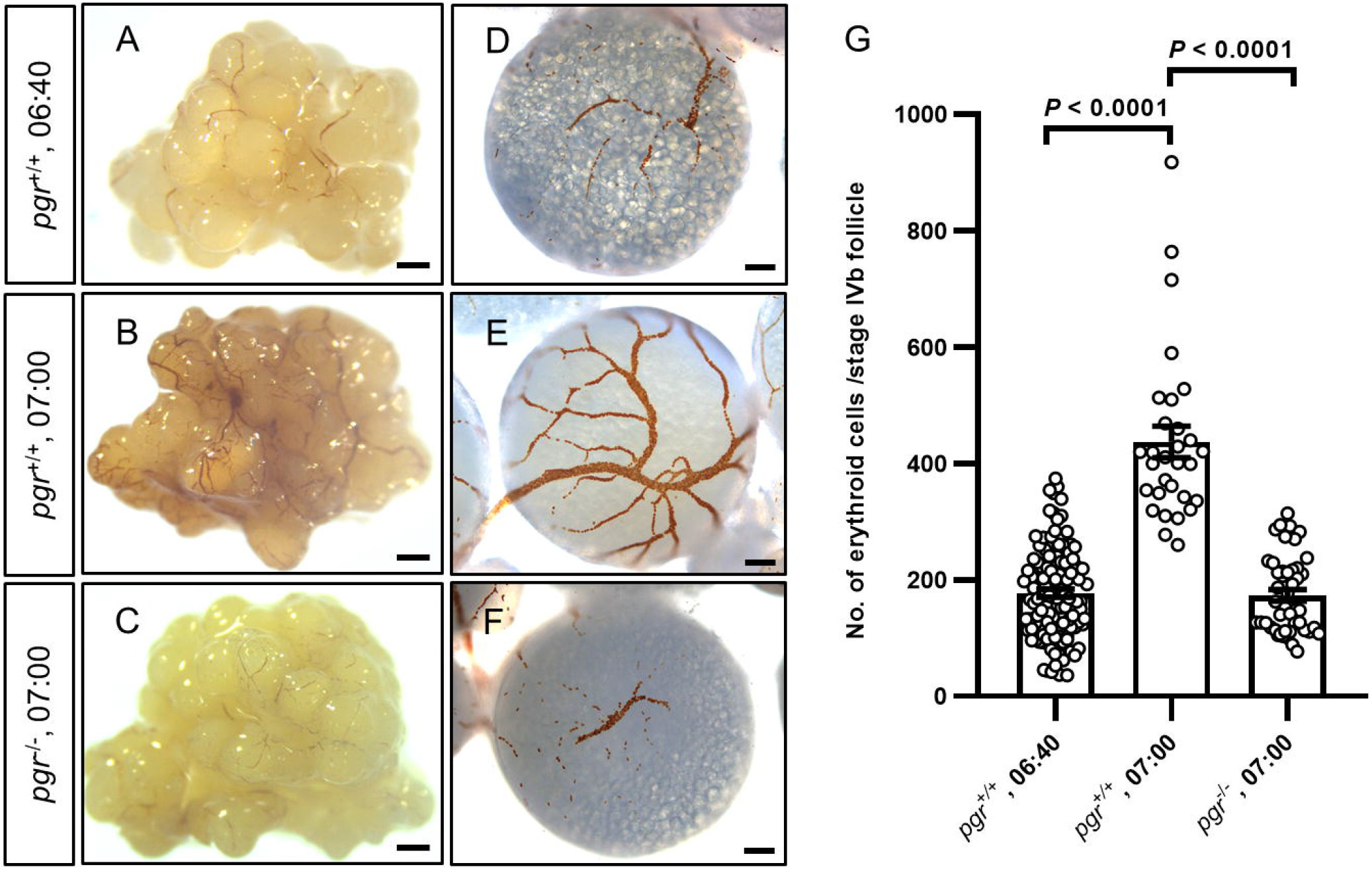
Increase of erythrocytes in capillaries on the surface of mature follicles during progression but prior to rupture. (**A, D, B, E**) Erythrocytes within the surface capillaries of mature follicles (IVb) from wildtype (*pgr*^*+/+*^) female zebrafish collected at 6:40 or 7:00 were visualized using *o*-dianisidine staining as follicles progressing towards rupture and ovulation. (**C, F**) Erythrocytes were also visualized in similar mature follicles that were collected at 7:00 from *pgr*^*-/-*^ females, in which ovulation never occurs (13). (**G**) At least 30 mature follicles were randomly chosen from at least 3 different females of each condition, and the number of erythrocytes in capillaries was counted by a blinded observer. Data are expressed as mean ± SEM (N ≥ 30). P-values were calculated by two-tailed Student’s t-test. Scale bars: 500 μm (A-C) or 100 μm (D-F).

#### 3.3d Inhibition of ovulation by anticoagulants *in vitro*

Intact IVb or ovulated follicles from *Tg(fli1:eGFP)* were easily recognizable by the interconnecting blood vessels with strong GFP expression in vascular endothelial cells and dramatic changes in size (Fig. 8A-D). As expected, ovulation was inhibited when follicles were exposed to EDTA (2 mM), an inhibitor of metalloproteases that are required for ovulation (38). Importantly, all four anticoagulants tested, i.e., heparin (100 µg/mL), warfarin (50 µg/mL), dabigatran etexilate (50 µM), or rivaroxaban (250 µM) significantly reduced the rates of ovulation (Fig. 8E-H).

**Figure 8.**
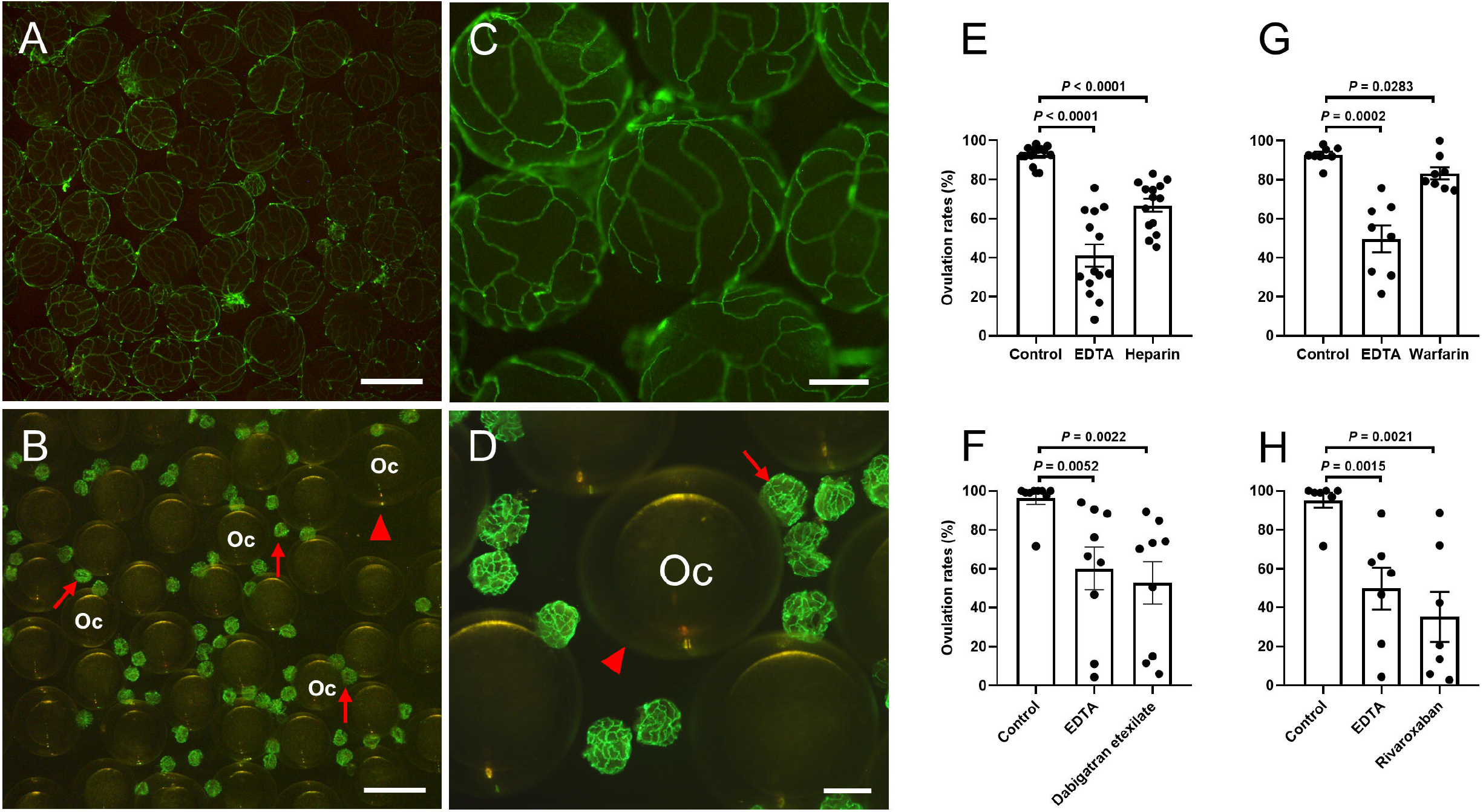
Inhibition of ovulation by anticoagulants *in vitro*. Ovulation rate was determined using mature follicles (IVb) isolated from transgenic female zebrafish (*Tg(fli1:eGFP*). The intact IVb follicles were easily recognizable by the interconnecting blood vessels with strong GFP expression in vascular endothelial cells. (**A, C**). After ovulation, oocytes were easily recognizable by the absence of interconnecting GFP-positive vessels and expansion of the vitelline envelope (chorion) (**B, D**). Follicular layers separated from ovulated oocytes could also be easily distinguished due to interconnecting blood vessels and a dramatic size reduction (**B, D**). (**A, C**) Mature follicles (IVb) were collected from *Tg(fli1:eGFP)* female zebrafish. (**B, D**) Representative images of ovulated mature follicles were taken 2 hrs after incubation at 28°C. Arrows indi cate the remaining follicular layer following ovulation. Arrowheads indicate expanded vitelline envelopes. Oc, oocyte. (**E-H**) The rates of ovulation *in vitro* were determined following 2 hrs incubation of mature follicles in a medium containing a vehicle control, EDTA (2 mM), or an anticoagulant (heparin,100 µg/mL; warfarin 50 µg/mL ; dabigatran etexilate, 50 µM; or rivaroxaban, 250 µM). Data are expressed as the mean ± SEM (N ≥ 7). P-values were calculated by paired two-tailed Student’s t-test. Scale bars: 1 mm (A, B) or 250 μm (C, D).

#### 3.3e Inhibition of ovulation by an anticoagulant *in vivo*

After warfarin treatment, none of the females showed evidence of ovulation, as determined by egg-laying *in vivo* (Fig. 9A) and examination of dissected ovaries (Fig. 9C, E). In contrast, females from the control group demonstrated high rates of ovulation (Fig. 9A, B, D). In control females, the numbers of erythrocytes in capillaries of preovulatory follicles (IVb) significantly increased prior to ovulation, an indicator of possible blood clotting prior to ovulation, as demonstrated in section 3.3c and figure 7. In contrast, few erythrocytes in capillaries of similar mature follicles (IVb) were observed in warfarin-treated female zebrafish, an indicator of normal blood flow in the vessels of follicles in these control fish (Fig. 9F-J).

**Figure 9.**
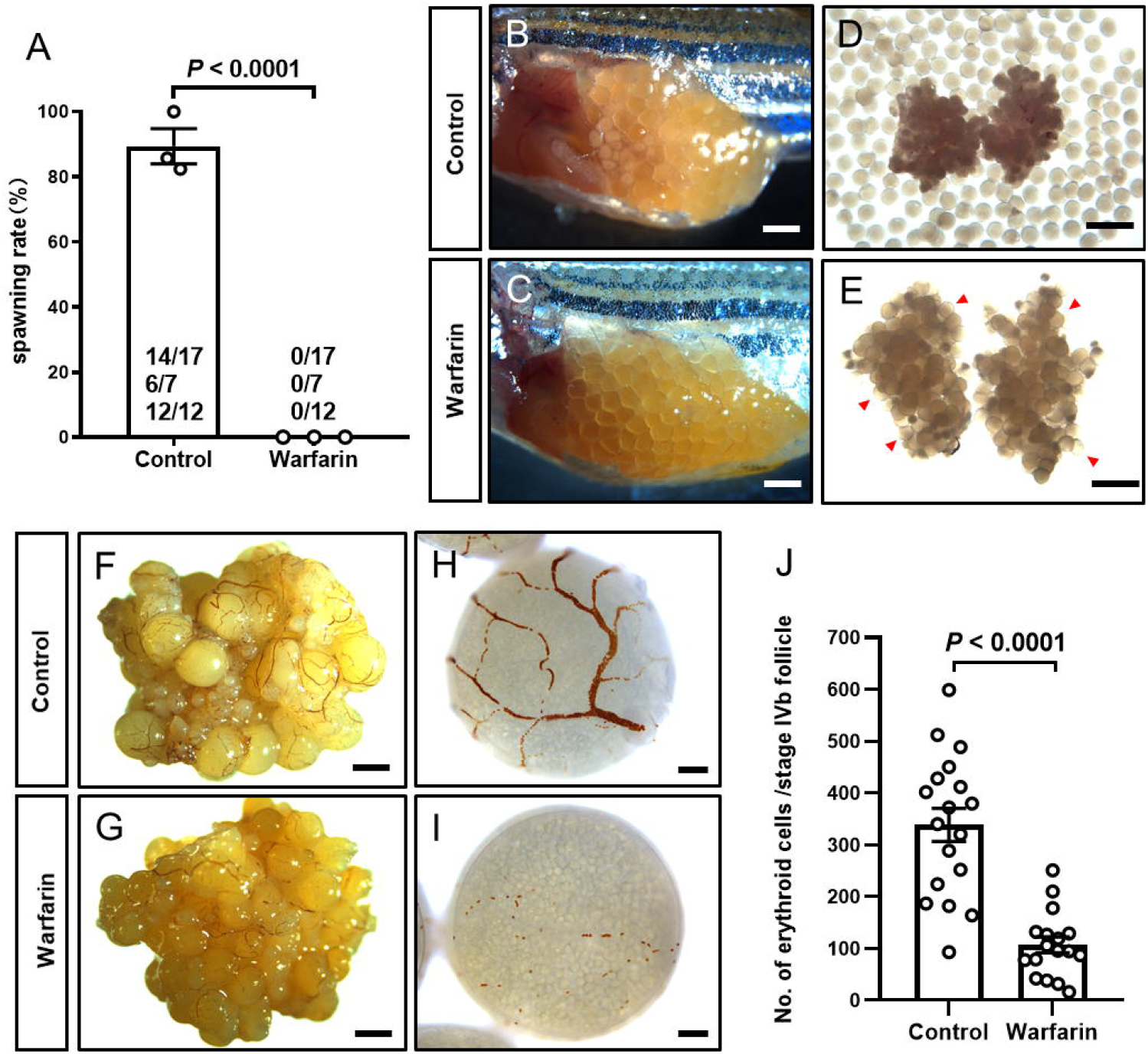
Inhibition of ovulation by an anticoagulant *in vivo*. (**A**) Showing average spawning rates from three repeated experiments in control (no exposure) or warfarin exposed female zebrafish. None of the warfarin-treated females laid eggs, as monitored by a blinded examiner, while control females had high rates of egg-laying (ovulation). The spawned/total number of females used in the first, second, and third experiments are listed inside the bar graph. (**B, D**) A representative ovary and ovulated eggs in the ovaries collected from control females. (**C, E**) Anovulation with all mature oocytes trapped within the ovary of warfarin-treated females. Arrowheads indicate anovulatory oocytes. (**F-I**) An increase of erythrocytes in the capillaries of mature follicles (IVb) collected from control fish prior to ovulation, while the number of erythrocytes in mature follicles of warfarin-treated females was dramatically less. (**J**) Inhibition of aggregation of red blood cells in warfarin treated preovulatory follicles in zebrafish. Data are expressed as the mean ± SEM (N ≥ 17). P-values were calculated by two-tailed Students’ t-test. Scale bars: 1 mm (B-E, F, G) or 100 μm (H, I).

#### 3.3f Reduced fertility in *f5*^*+/-*^ zebrafish females *in vivo*

Although *f5* homozygous mutants live to early adulthood, most die by 4-6 months of age and do not breed well (45). In contrast, *f5*^+/-^ zebrafish are indistinguishable from wildtype siblings, and no abnormal development of any organs have been observed (45). Therefore, we chose to use *f5*^+/-^ for the examination of female fertility in a consecutive spawning test. We observed significantly reduced numbers of embryos per spawning and total embryos over two weeks in comparison to wildtype sibling females. (Fig. 10A, B). The spawning frequency of *f5*^*+/-*^ female fish was also significantly decreased in comparison to wildtype siblings (Fig. 10C). One *f5*^*+/-*^ female only spawned once in two weeks (Fig. 10C), and the spawning intervals of *f5*^*+/-*^ females were significantly longer in comparison to that of wildtype siblings (Fig. 10D).

**Figure 10.**
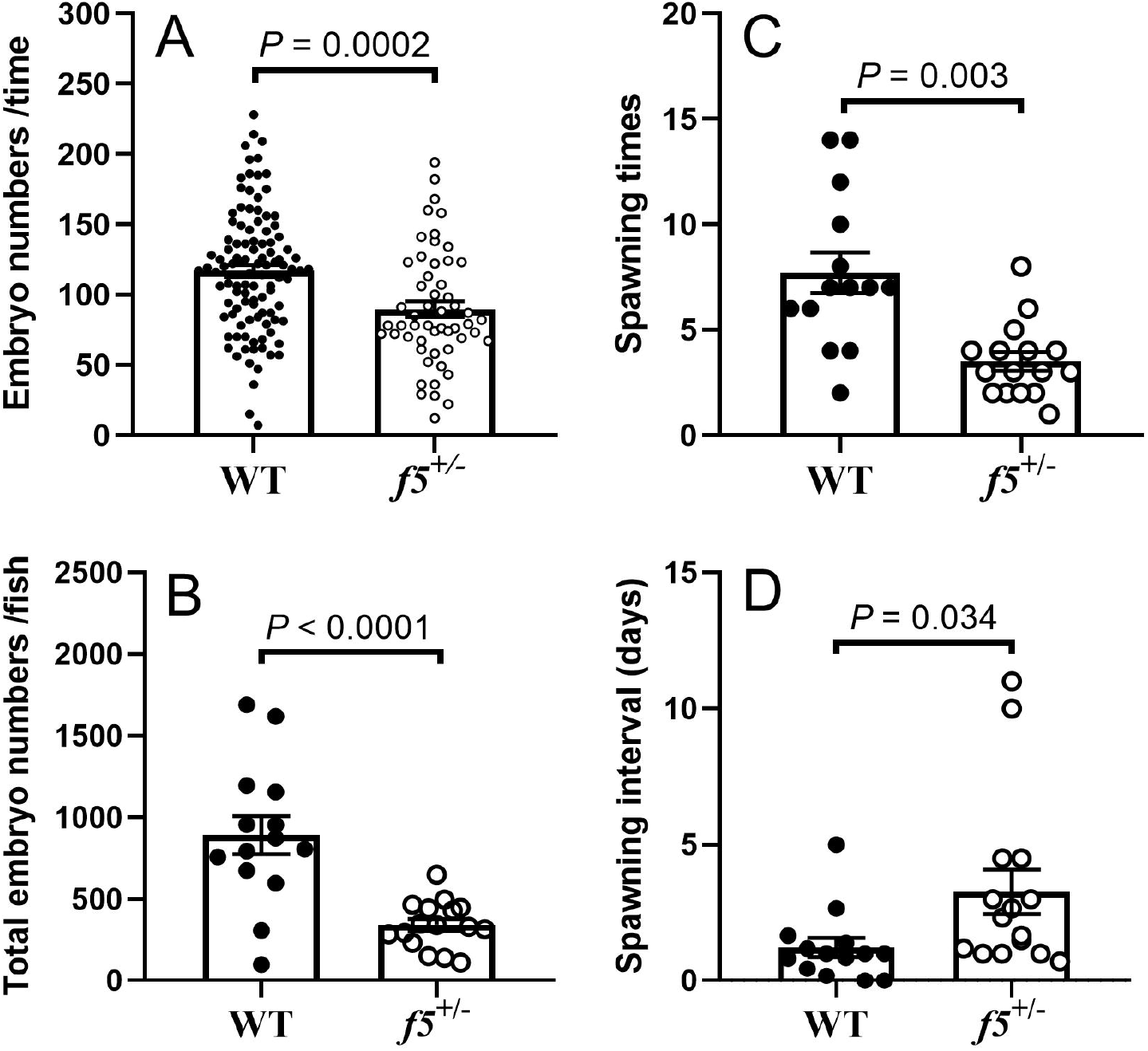
Reduced fecundity in *f5*^*+/-*^ female zebrafish. (**A**) Reduced numbers of live embryos per spawning for each *f5*^*+/-*^ female in comparison to that of wildtype (*f5*^*+/+*^) females. (**B**) Reduced total numbers of embryos for each *f5*^*+/-*^ female fish in comparison to wildtype (*f5*^*+/+*^) females. (**C**) Reduced spawning times in *f5*^*+/-*^ females in comparison to that of wildtype (*f5*^*+/+*^) females. (**D**) Increased spawning intervals in *f5*^*+/-*^ females in comparison to wildtype (*f5*^*+/+*^) females. Data are expressed as the mean ± SEM (N ≥ 14). P-values were calculated by two-tailed Student’s t-test.

## 4. Discussion

We have provided comprehensive evidence demonstrating that the expression of coagulation factors *f5* and *f3a* in preovulatory follicles is controlled by progestin and its cognate receptor Pgr. Increased expression of these coagulation factors is likely critical for the regulation of ovulation and female fertility in addition to their well-established roles in hemostasis. Our conclusions are based on the following evidence: 1) Coagulation factors, *f5* and *f3a*, are expressed in the follicular cells and increase dramatically prior to ovulation during natural spawning cycles in zebrafish; 2) Expression of these cofactors in preovulatory follicles is controlled by progestin via Pgr, which are essential and upstream regulators for ovulation; 3) Follicular cells with strong Pgr expression are adjacent to blood capillaries; 4). As follicular cells near rupture, a significant increase in the numbers of erythrocytes in blood vessels suggests blood clotting which is correlated with a dramatic increase of *f5* and *f3a* in preovulatory follicular cells; 5) Inhibition of coagulation leads to reduced ovulation; 6) Low fertility is found in *f5*^*+/-*^ zebrafish female mutants.

A major finding of the present study is the regulation of *f5* and *f3a* by progestin via Pgr in the preovulatory follicles during ovulation (right side of Fig.11). F3 is expressed in a wide array of tissues (46). It has been demonstrated that physiological concentrations of P4 could increase both F3 mRNA and protein in breast cancer cell lines ZR-75 and T47D (47). Increased F3 expression was correlated with an increase of procoagulant activity, which was hypothesized to be responsible for the increased risk of thrombosis in breast cancer patients (48). Studies of the uterus have also shown that RU486, an antagonist of PGR, not only blocks but also reverses progestin-enhanced expression of F3 mRNA and protein in stromal cells (49). In line with these studies, PGR binding sites were identified in the promoter region of the human F3 gene (28,50). The increase of *f3a* expression in the preovulatory follicles is primarily controlled by Pgr (current study). A difference in the weak activity of *f3a* promoters observed in a human immortal cell line (HEK293T) *in vitro* in comparison to progestin-induced strong *f3a* promoter activities found in zebrafish follicles both *in vivo* and *in vitro* may be due to missing core *cis*-regulatory elements in the cloned promoter sequence of zebrafish *f3a* (28). Nevertheless, our results demonstrate that like *f5, f3a* expression is also controlled by progestin via Pgr (Fig.11) (28).

Until now, no information has been available regarding progestin regulation of F5. Estrogen and estrogen receptors regulate *F5* expression in mouse bone marrow/bone (51,52). The present study demonstrates that a new extrahepatic site of *f5* expression is regulated by progestin through Pgr in zebrafish. Moreover, progestin enhances the promoter activities of *f5* in zebrafish and F5 in humans via PGR. Several putative PREs were predicted in the promoter regions of human, mouse, and zebrafish *F5* (Supplemental Table 5) (28). Our site-directed mutation analyses of these PREs in the zebrafish *f5* promoter suggest direct regulation of progestin/Pgr via binding to and activating these PREs. One PRE (5’-TGACTCA-3’) in the *f5* promoter is also conserved between zebrafish, mice and humans (Supplemental Table 5) (28). Whether P4 and/or PGR regulate *F5* expression in human ovaries *in vivo* will be an interesting topic for further investigation.

The functions of coagulation factors in hemostasis are well established (53-55). F3 is a cell surface glycoprotein that is a potent initiator for activation of procoagulant proteases in the clotting cascade. Activated F5 serves as a cofactor protein for the activated serine protease factor Xa in the prothrombinase complex, which rapidly converts prothrombin to thrombin. Several lines of evidence including biochemical and evolutionary analyses suggest that the coagulation system of zebrafish is nearly identical to mammals (19-21). However, the role of coagulation factors during ovulation is significantly less studied. The formation of blood clots at apical blood vessels was found shortly before follicular rupture in several mammalian species (1,3-6). In addition, the permeability of blood vessels increased 3-fold prior to ovulation in rats (56), which likely facilitates material exchange between the follicles and blood vessels. The increase of material exchange might be important for anticipating local but avoiding systemic activation of blood clotting. Our observation of ruptured blood vessels on the surface of ovulating follicles further supports this hypothesis. In addition, we also found a significant increase of erythrocytes in capillaries on the surface of ovulating follicles as they progress to rupture/ovulation, suggestive of blood clotting, ruptured blood vessels, and/or vasoconstriction (57-61). Lack of functional F5 or F8 causes various bleeding symptoms, including recurrent and serious ovulation-related blood loss in women (62). The occurrence of blood clotting in advance of follicular rupture could preemptively stop excessive bleeding caused by ovulation-induced capillary rupture. The granulosa cells appear to be the primary cells for synthesizing F5 and F3a, as these account for most that express Pgr (this study & (10,41)). Intriguingly, we also found that strong Pgr expressing theca cells were localized near capillaries on the follicles. Whether these theca cells are key switchers of F3a-initiated hemostasis in the capillaries around ovarian follicles will be an interesting topic to examine in the future.

The dramatic induction of *f5* and *f3a* expression in preovulatory follicles via Pgr, an essential mediator for activation of various ovulation-related genes, suggests that locally produced F5 and F3a may be required for ovulation. Activation of multiple metalloproteases within the ovulating follicles at the right times is a critical step for sequential digestion of extracellular matrix proteins of the follicular cells that are required for ovulation (1,42). Their activation and substrate binding are precisely regulated by the timing of prodomain cleavage of the enzyme and removal of endogenous tissue inhibitors of metalloproteases (Timps) (63). Protease inhibitors (Timps and other inhibitors) are typically present at high concentrations in the serum to prevent metalloproteases from acting accidentally (64, left side of Fig.11). Vasoconstriction of blood vessels within the thecal layer at the apex of the follicles was suggested to be critical for the removal of protease inhibition from the circulation (65). We hypothesize that coagulation that occurs shortly before ovulation in the follicles may have a similar function. In addition to the removal of protease inhibitors from serum, the cleavage of propeptide from latent form of metalloproteinase is required for activation and thus ovulation (Fig.11). An F3-dependent pathway of thrombin generation was also shown in bovine, equine, and human follicular fluid (7,66,67). Granulosa cells were shown to express and secrete fibrinogen, which is catalyzed by thrombin to become fibrin (68). Fibrin formation could stimulate activities of plasmin/plasminogen activators (plat and plau) signal, which is well-established for plasminogen (plg) activation (69). Furthermore, the function of Plat, Plau, and Plg in metalloproteinase activations during ovulation is also recognized (70,71). Therefore, we hypothesize that a dramatic increase of F3a and F5 expression in the follicular cells may facilitate the formation of fibrin, followed by the activation of Plat, Plau, and Plg, which further activate metalloproteinases. Taken together, as blood coagulation occurs, blood flow decreases, then stops completely, effectively terminating inhibitory effects of protease inhibitors from the serum, triggering metalloprotease activations to start the degradation of the follicular extracellular matrix for preparing the release of a mature oocyte (right side of Fig.11). Consistent with findings from medaka (38), the rate of ovulation was suppressed by *in vitro* EDTA treatment in the current study likely via inhibition of metalloproteases, or possibly inhibition of coagulation. EDTA is a known anticoagulant, acting through chelation of calcium, as many coagulation factors depend on this ion for function. We purposely selected warfarin over three other anticoagulants because of its well-established effects *in vivo*. As we hypothesized, four days of *in vivo* exposure to warfarin completely suppressed (100%) ovulation in female zebrafish.

Another finding in the present study is that reduced fecundity and impaired ovulation were observed in *f5*^*+/-*^ female zebrafish *in vivo*. In humans, a patient diagnosed with severe F5 deficiency had to undergo bilateral ovariectomies because of the presence of hemorrhagic anovulatory follicles and life-threatening hemorrhage in the pouch of Douglas (72). Circulating F3 has been shown to be elevated in women with polycystic ovarian syndrome, the most common form of anovulatory infertility (9). Together, these observations argue for conserved functions of coagulation factors in the regulation of ovulation in vertebrate ovaries. Long-term efforts in the development of new instruments and new techniques/methods, with suitable *in vivo* imaging and examining the association of blood flow and timing of metalloproteases activation, are required for further elucidation of underlying mechanisms and testing our hypothetical model (Fig.11).

In summary, our results demonstrate that *f5* and *f3a* expression are dramatically induced by progestin via a Pgr-dependent mechanism in preovulatory follicles (Fig. 11). Moreover, these coagulation factors may regulate ovulation and female fertility in addition to their roles in hemostasis. Future studies on the regulation and functions of coagulation factors in other organisms including mammalian follicles or ovarian fluid during ovulation will likely lead to developments of diagnostic and therapeutic modalities for patients with gynecological problems associated with mutations and/or dysregulation of coagulation.

**Figure 11.**
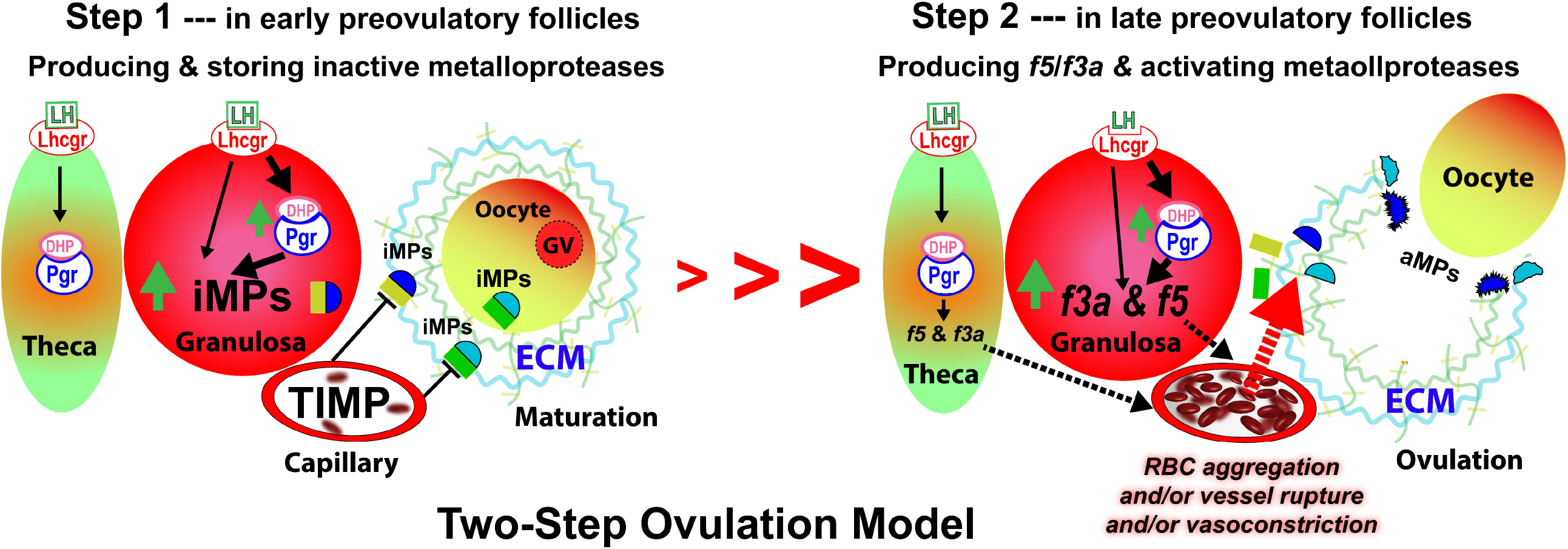
A proposed two-step ovulation model for producing and activating metalloproteases via nuclear progestin receptor (Pgr) mediated upregulation of coagulation factor *f5* and *f3a* in ovarian follicular cells in zebrafish. It is well established that luteinizing hormone (LH) induces ovulation via LH receptor (Lhcgr) in preovulatory follicles, namely in stage IV follicles in zebrafish. LH signaling is primainly mediated by progestin (DHP, i.e., 17α,20β-dihydroxy-progesterone, a native ovulation-inducing steroid in zebrafish) and its receptor (Pgr) in the production and activation of metalloproteases. However, how metalloproteases are activated during ovulation is still unclear.

## Acknowledgments

We want to thank Dr. Mingyu Li (Xiamen University, China) and Dr. Yu Xue (Minnan Normal University, China) for providing zebrafish transgenic lines (*Tg(fli1:eGFP), Tg(fli1:DsRed)*), Dr. Shuang-Bo Kong (Xiamen University, Xiamen, China) for sharing human PGRB expression vector (pCMV-HA-hPGRB). This work was supported by Fundamental Research Funds for the Central Universities (No. 20720200114), NSFC 31672628, and NSFC 41976092 to SXC, NIH GM100461 to YZ, and NIH R35 HL150784 to JAS.

## Data Availability Statement(s)

All data generated or analyzed during this study are included in this published article or in the data repositories listed in References 28.

## Facts

The illustration on the left side is shows the upregulation of metalloproteases (blue dome activation domains) linked to yellow or green square (propeptide inhibitory domain), see ref# 63 for detail) in early preovulatory follicles, to be more specific, during the transition from stage IVa to early IVb follicles, which is an important preparation step for ovulation (16,29,30). However, these proteases are produced in a latent form, i.e., inactive form of metalloproteases (iMPs, see ref# 63 for detail), and inhibited by propeptide domain and serum inhibitors such as tissue inhibitors of metalloproteases (TIMP) presented at high levels in and delivered by blood capillaries to the follicles (73-77). The illustration on the right side shows a dramatic increase of coagulation factors *f5* and *f3a* shortly before ovulation in the late stage of preovulatory follicles (late IVb follicles in the case of zebrafish), which are the key step for ovulation and also mainly upregulated by progestin via Pgr (current study).

## Main hypothesis

In this study, we observed increased aggregation of red blood cells (RBC) in capillaries located on the surface of preovulatory follicles, which was likely due to: 1) Occurrence of blood clotting, and/or 2) Ruptured blood vessels (see figure 6 and supplemental video clip A), and/or 3) vasoconstriction (another possibility not examined in the current study). These ovarian follicles derived, and locally produced coagulation factors (F5 and F3a) are imported into inside blood vessels meshed in the follicles and activated via yet to be defined mechanisms. Once F5 and F3a are activated, these coagulation factors accelerate blood coagulation, which in turn stops blood flow and therefore effectively terminates serum inhibition including TIMP inhibition of metalloproteases from the bloodstream. In addition, the production of fibrin (not shown) due to the formation of blood clots stimulates activities of plasmin/plasminogen activators, which are well-known inducers for metalloprotease activation. The combined force of these two pathways triggers rapid activation of metalloproteases, breakdown of extracellular matrix and basal membrane, rupture of follicular cell layers, and the final release of a mature oocyte.

## lternative Hypothesis

Vasoconstriction and/or ruptured blood vessels also could be the cause leading to local erythrocytes aggregation and effectively reduces blood flow and local Timp contents, activating metalloproteases via similar mechanisms described above.

### Abbreviations

iMPs: inactive metalloproteases
aMPs: activated metalloproteases
RBC: red blood cell
LH: luteinizing hormone
Lhcgr: luteinizing hormone/choriogonadotropin receptor
DHP: 17α, 20β-dihydroxy-4-pregnen-3-one
Pgr: nuclear progestin receptor:
ECM: extracellular matrix
GV: germinal vesicle
TIMP: tissue inhibitor of metalloproteases
Va: stage IVa ovarian follicle, fully-grown but immature follicle
IVb: stage IVb follicle, mature but yet ovulated follicle
f5: coagulation factor f5
f3a: coagulation factor 3a, also named as tissue factor (TF).

