## Supplemental information for "Nuclear Progestin Receptor Mediated Linkage of Blood Coagulation and Ovulation"

**SUPPLEMENTAL MATERIALS**

**Supplemental Table 1 |** Zebrafish knockouts and transgenic lines used.

| Knockouts lines | Reference |
| --- | --- |
| *pgr^-/-^*  *f5^+/-^* | (1)  (2) |
| Transgenic lines | **Reference** |
| *Tg(pgr:eGFP)*  *Tg(fli1:eGFP)*  *Tg(**fli1:DsRed)*  *Tg(pgr:eGFP/ fli1:DsRed)* | (3)  (4)  (4)  this study |

1.	Liu DT, Carter NJ, Wu XJ, Hong WS, Chen SX, Zhu Y. Progestin and Nuclear Progestin Receptor Are Essential for Upregulation of Metalloproteinase in Zebrafish Preovulatory Follicles. *Frontiers in Endocrinology*. 2018;9.

2.	Weyand AC, Grzegorski SJ, Rost MS, Lavik KI, Ferguson AC, Menegatti M, Richter CE, Asselta R, Duga S, Peyvandi F, Shavit JA. Analysis of factor V in zebrafish demonstrates minimal levels needed for early hemostasis. *Blood Adv*. 2019;3(11):1670-1680.

3.	Huang J, Zhang TT, Jiang K, Hong WS, Chen SX. GFP expression pattern in pituitary and gonads under the control of nuclear progesterone receptor promoter in transgenic zebrafish. *Developmental Dynamics*. 2020.

4.	Lawson ND, Weinstein BM. In vivo imaging of embryonic vascular development using transgenic zebrafish. *Dev Biol*. 2002;248(2):307-318.

**Supplemental Table 2 |** Sources of major chemicals and reagents.

| Name | Vendor | Catalog Number |
| --- | --- | --- |
| 17α, 20β-dihydroxy-4-pregnen-3-one (DHP)  Progesterone (P4)  EDTA  Heparin  Human chorionic gonadotropin (hCG)  Leibovitz's L-15 medium  Paraformaldehyde  Hochest 33342  *O*-dianisidine  Warfarin | Steraloids  Steraloids  Sinopharm Group Co. Ltd.  Sinopharm Group Co. Ltd.  ProSpecTany TechnoGene Ltd.  Thermo Fisher Scientific  Merck  Beyotime Institute of Biotechnology  Macklin  Sigma-Aldrich | Q1850-000  Q2600-000  10009617  63007131  HOR-250  21083027  8187151000  C1029  D807428  A4571 |
| Dabigatran etexilate | Apexbio | A8381 |
| Rivaroxaban | Apexbio | A4338 |

**Supplemental Table 3 |** Oligo primers used.

| Gene | Forward primer (5′→3′) | Reverse primer (5′→3′) | Annealing [temperature](https://cn.bing.com/dict/clientsearch?mkt=zh-CN&setLang=zh&form=BDVEHC&ClientVer=BDDTV3.5.1.4320&q=%E9%80%80%E7%81%AB%E6%B8%A9%E5%BA%A6" \t "_blank) | Accession number |
| --- | --- | --- | --- | --- |
| For qPCR  *f3a*  *f5*  *ef1a* | GCTTGTATTGGCGCTGGTTT  ACGGCTACACAAATGGCTCA  GCGCAAGGAGGGTAATGCTA | TCGTCCTGATGCAGTATGGC  GGTGGACACCGGTCATACTG  GGGCGAAGGTCACAACCATA | 60 ℃  60 ℃  60 ℃ | NM_001245967  NM_001328531  NM_131263 |
| For RT-PCR  *f3a*  *f5*  *ef1a* | GCTTGTATTGGCGCTGGTTT  GTCCTCAGGGTTACTTGGGC  GCGCAAGGAGGGTAATGCTA | CAGACTAAAGAACCCACC  TGGCGCCGTATACTCATCAC  GGGCGAAGGTCACAACCATA | 58 ℃  58 ℃  58 ℃ | NM_001245967  NM_001328531  NM_131263 |
| For WISH  *f3a*  *f5* | GCTTGTATTGGCGCTGGTTT  GTCCTCAGGGTTACTTGGGC | CAGACTAAAGAACCCACC  TGGCGCCGTATACTCATCAC |  | NM_001245967  NM_001328531 |
| For luciferase assay  *pgr*  *f3a*  *f5*  *F5** | CTCGCTAGCATGGACACGGTGAA  CGCGCTAGCACAAATCATGAGCCT  CGTGCTAGCTGTCTGCTGAAGTATG  CCAGCTAGCGCACACCCTACACTGC | CCGAAGCTTTCATTTGTGGTGAA  TTTCCATGGATTGTCCCAAGTCCA  GATCTCGAGGTGAAGAGAAGTATGA  CCGCCATGGGCTTCCTTTCCTGCTCCC |  | NM_001166335  ENSDARG00000099124  ENSDARG00000055705  ENSG00000198734 |

* indicated the gene of human.

**Supplemental Table 4 |** Plasmid constructs for Pgr expression and dual luciferase assay.

| Plasmid name (gene/reporter) | Backbone | Insert | [Species](https://cn.bing.com/dict/clientsearch?mkt=zh-CN&setLang=zh&form=BDVEHC&ClientVer=BDDTV3.5.1.4320&q=%E7%89%A9%E7%A7%8D) |
| --- | --- | --- | --- |
| pcDNA3.1(+)-zPgr (Pgr)  pCMV-HA-hPGRB (PGRB)  pGL3-zf3a (firefly luciferase)  pGL3-zf5 (firefly luciferase)  pGL3-hF5 (firefly luciferase)  pGL3-MMTV (firefly luciferase)  pRL-TK (*Renilla* luciferase) | pcDNA3.1(+)  pCMV-HA  pGL3  pGL3  pGL3  pGL3  pRL | *pgr* cDS  *PGRB* cDS  *f3a* promoter (-1892/+133)  *f5* promoter (-2063/+32)  *F5* promoter (-2125/+202)  mouse mammary tumor virus promoter  herpes simplex virus thymidine kinase promoter | zebrafish  human  zebrafish  zebrafish  human |

**Supplemental Table 5 |** Predicted PGR binding sites in the proximal promoter sequence of human, mouse, and zebrafish *F5* gene.

| Promoter of Gene | TF | Source* | Start | Stop | Strand | Matched Sequence |
| --- | --- | --- | --- | --- | --- | --- |
| Human *F5* | PGR  PGR  PGR  PGR  PGR  PGR  PGR  PGR  PGR  PGR  PGR  PGR | hTFtarget*  hTFtarget  hTFtarget  hTFtarget  hTFtarget  hTFtarget  hTFtarget  hTFtarget  hTFtarget  database**  database  hTFtarget | -1878  -1676  -1193  -2339  -1065  -1328  192  -1681  -1324  -1715  -1081  -1113 | -1867  -1669  -1186  -2330  -1056  -1319  201  -1672  -1312  -1702  -1068  -1097 | -  -  +  -  +  +  +  -  -  -  -  + | TGTTTGCTTAT  TGACTCA  CAGGACA  GGCAGGAAG  GACAGGAAA  GGGAGGAAG  GAAAGGAAG  CTCAGGAAA  ACAGCCTCTTCC  AGAACACCTAGCC  AGGACCCCTTGTA  GAAACAGTCAGATCCT |
| Mouse *F5* | PGR  PGR  PGR  PGR  PGR PGR  PGR  PGR  PGR  PGR PGR  PGR  PGR  PGR  PGR PGR  PGR  PGR  PGR  PGR PGR  PGR | hTFtarget  database  hTFtarget  hTFtarget  hTFtarget  hTFtarget  hTFtarget  hTFtarget  hTFtarget  hTFtarget  hTFtarget  hTFtarget  hTFtarget  hTFtarget  hTFtarget  database  database  database  database  hTFtarget  hTFtarget  hTFtarget | -132  -615  -995  -386  112  -614  -127  -1951  -2665  -2409  -2581  -2543  -2614  -2576  -2538  -1892  -1878  -1892  -1878  -822  -2619  -2088 | -116  -602  -988  -379  119  -602  -118  -1942  -2656  -2400  -2565  -2527  -2605  -2567  -2529  -1885  -1871  -1885  -1871  -810  -2603  -2072 | -  -  +  -  +  +  -  +  +  -  -  -  -  -  -  -  +  +  -  +  -  - | GGGAAAGGAAGGGCAA  GGGACAAACTGTT  TGACTCA  CAGGACA  CAGGACA  ACAGTTTGTCCC  GAAAGGAAG  CAGAGGAAA  GACAGGAAC  GTGAGGAAG  GGAGCAGGAAGGGGCG  GGAGCAGGAAGGGGCG  AGCAGGAAG  AGCAGGAAG  AGCAGGAAG  AGAACAG  AGAACAG  CTGTTCT  CTGTTCT  ACTGGATGTGCT  GGAGCAGGAAGGGGTG  AAAACACAAAGAGCTT |
| Zebrafish *f5* | PGR  PGR  PGR  PGR | hTFtarget  database  hTFtarget  hTFtarget | -863  -1438  -1179  -964 | -856  -1425  -1170  -948 | -  +  -  +  +  + | TGACTCA  TGAGTCA  AGGACATTTTTTA  TAAAAAATGTCCT  CAGAGGAAA  TGAACAATTAGTCCTG |

The proximal promoter sequence (-2601/+202) of the human *F5* gene (ENSG00000198734), (-2712/+125) of the mouse *F5* gene (ENSMUSG00000026579), and (-2063/+32) of the zebrafish *f5* gene (ENSDARG00000055705) was obtained from ENSEMBL genome browser (https://asia.ensembl.org/index.html). The PGR binding sites were predicted using human transcription factor database (http://bioinfo.life.hust.edu.cn/HumanTFDB/#!/tfbs_predict).

* hTFtarget means transcription factor bind site was predicted based on the motifs collected from Database of Human Transcription Factor Targets (http://bioinfo.life.hust.edu.cn/hTFtarget#!/).

** database means the transcription factor bind site was predicted based on the motifs collected from TRANSFAC, JASPAR, CIS-BP, and HOCOMOCO.

**Supplemental Table 6 |** Predicted PGR binding sites in the proximal promoter sequence of human and mouse *F3* gene, and zebrafish *f3a* gene.

| Promoter of Gene | TF | Source | Start | Stop | Strand | Matched Sequence |
| --- | --- | --- | --- | --- | --- | --- |
| Human *F3* | PGR  PGR  PGR  PGR  PGR | hTFtarget*  database**  hTFtarget  database  hTFtarget | -1512  -2087  -1828  -1613  -1613 | -1505  -2074  -1819  -1600  -1597 | -  -  +  +  + | TGACTCA  AGGACATTTTTTA  CAGAGGAAA  TGAACAATTAGTC  TGAACAATTAGTCCTG |
| Mouse *F3* | PGR  PGR  PGR  PGR  PGR PGR  PGR  PGR  PGR  PGR PGR  PGR  PGR  PGR  PGR PGR  PGR  PGR  PGR  PGR PGR | database  hTFtarget  hTFtarget  hTFtarget  hTFtarget  hTFtarget  hTFtarget  hTFtarget  hTFtarget  hTFtarget  hTFtarget  hTFtarget  hTFtarget  hTFtarget  database  database  database  database  hTFtarget  hTFtarget  hTFtarget | -439  -2517  -714  -2540  -135  -2171  -1589  -1175  -1107  -319  -1590  -1478  -1299  -2518  -2166  -856  -2166  -856  -2518  -1589  -2518 | -426  -2501  -703  -2533  -128  -2159  -1573  -1166  -1096  -310  -1574  -1469  -1290  -2502  -2159  -849  -2159  -849  -2502  -1573  -2502 | -  +  +  -  -  +  +  +  +  +  -  -  +  +  -  +  +  -  -  +  - | AGATCATTCTGTC  AGAACTCTGCGTTCTC  TGTTTATTTTT  TGACTCA  TGACTCA  TCATTCTGTTCT  TGAACAATAAGTTCCC  GAAAGGAAG  TATTTACATTA  CAGAGGAAA  GGAACTTATTGTTCAT  CAAAGGAAG  AGCAGGAAG  CAGAACTCTGCGTTCT  AGAACAG  AGAACAG  CTGTTCT  CTGTTCT  AGAACGCAGAGTTCTG  TGAACAATAAGTTCCC  AGAACGCAGAGTTCTG |
| Zebrafish *f3a* | PGR | hTFtarge | -1384 | -1373 | - | TGTTTATTTTT |

The proximal promoter sequence (-2712/+123) of the human *F3* gene (ENSG00000117525), (-2640/+178) of the mouse *F3* gene (ENSMUSG00000028128), and (-1892/+133) of the zebrafish *f3a* gene (ENSDARG00000099124) was obtained from ENSEMBL genome browser (h`ttps://asia.ensembl.org/index.html). PGR binding sites were predicted using human transcription factor database (http://bioinfo.life.hust.edu.cn/HumanTFDB/#!/tfbs_predict).

* hTFtarget means transcription factor bind site was predicted based on the motifs collected from Database of Human Transcription Factor Targets (http://bioinfo.life.hust.edu.cn/hTFtarget#!/).

** database means the transcription factor bind site was predicted based on the motifs collected from TRANSFAC, JASPAR, CIS-BP, and HOCOMOCO.

**Supplement video clip A**

Illustration of capillary rupture and mature egg release by imaging a part of ovary undergoing ovulation process *in vitro* using a DFC 550 digital camera and an M165FC fluorescent dissecting microscope (Leica, Germany). This video clip was generated using a series of time-lapse images. The red arrow indicates a rupture of a blood vessel on the surface of a mature oocyte prior to its release during ovulation. Oc, oocyte.
